# A unique genomic island-governed cannibalism in *Bacillus* enhanced biofilm formation through a novel regulation mechanism

**DOI:** 10.1101/2021.10.04.462985

**Authors:** Rong Huang, Qing Li, Dandan Wang, Haichao Feng, Nan Zhang, Jiahui Shao, Qirong Shen, Zhihui Xu, Ruifu Zhang

## Abstract

Cannibalism is a differentiation strategy and social multicellular behavior in biofilms. The novel factors and mechanisms to trigger the bacterial cannibalism remain scarce. Here, we report a novel bacillunoic acids-mediated strategy for manipulating cannibalism in *Bacillus velezensis* SQR9 biofilm formation. A subfraction of cells differentiate into cannibals that secrete toxic bacillunoic acids to lyse a fraction of their sensitive siblings, and the released nutrients enhance biofilm formation. Meanwhile, the self-immunity of cannibal cells was induced by bacillunoic acids. A two-component system, the OmpS-OmpR signal-transduction pathway, controls the expression of the ABC transporter BnaAB for self-immunity. Specifically, bacillunoic acids activate the autophosphorylation of OmpS, a transmembrane histidine kinase, which then transfers a phosphoryl group to its response regulator OmpR. The phosphorylation of OmpR activates the transcription of the transporter gene *bnaAB* by binding its promoter. Thus, bacillunoic acids are pumped out of cells by the ABC transporter BnaAB. Moreover, we discovered that strain SQR9 could use the bacillunoic acids-mediated cannibalism to optimize its community to produce more bacillunoic acids for bacterial competition. This study revealed that bacillunoic acids play a previously undiscovered dual role in both cannibalism during biofilm formation and interspecies competition, which has an important biological significance.

## Introduction

Microorganisms in natural habitats usually exist as multicellular communities, such as biofilms, the predominant lifestyle of bacteria (1). Microbial cells encased in biofilms respond to different environmental signals and differentiate into distinct subpopulations (2). Under starvation, a subpopulation inside a bacterial biofilm autolyzes to sustain the survival of the remaining bacterial subpopulations, and this process has been shown in biofilms of both gram-positive and gram-negative bacteria (3-9). Bacterial autolysis is a coordinated cell death process and is defined as cell death in a subset of a bacterial population induced by other cells of the same species (10-11). Bacterial autolysis in biofilms provides nutrients for sibling cells, releases components of the biofilm matrix, such as DNA, and promotes the development of biofilms (12).

The control of bacterial cell death and lysis is an important mechanism in cell differentiation and development (3,5). Cannibalism, fratricide, and programmed cell death (PCD) are the three well-studied mechanisms to promote bacterial cell death. In *Bacillus subtilis*, once nutrients are insufficient, the low level of Spo0A-P triggers a subfraction of cells to differentiate into cannibal cells to secrete extracellular killing factors of the antimicrobial peptide SkfA or SdpC. Low level Spo0A-P can also activate the expression of the immunity gene *skfEF* in the same gene cluster with the toxin synthesis gene, while toxin SdpC binds immunity protein SdpI to subsequently sequester SdpR and allow *sdpIR* expression to protect the cannibal cells from the lysis of self-secreted toxins (13-15). These two cannibalism processes are controlled by two independent gene clusters: *skfA-H* and *sdpABC-RI*. Spo0A-inactive cells cannot express the *skf* and *sdp* clusters and hence are susceptible to the toxins produced by Spo0A-active cells (15-16). Lysed cells serve as nutrients to feed the cannibal subcommunity in biofilms (13,17).

In *Streptococcus pneumoniae*, competent cells produce peptides that target noncompetent siblings in a process known as fratricide. The rapid increase in cell density results in extracellular accumulation of competence-stimulating peptide; once exceeding a threshold concentration, it activates the *comDE* two-component system. This system regulates approximately 20 early competence (*com*) genes, including *comX*, which is required for the expression of cell lysis-related genes, and the immunity protein gene *comM*, which protects the competent subpopulation against autolysis (18-20). The expression of cell lysis-related genes encoding hydrolases *cbpD*, autolysin *lytA* and bacteriocin *cibAB* depends on ComX (10,21-23). The immune protein CibC is also coexpressed with CibAB to protect ComX-dependent competent cells (10). ComX-independent cells are autolyzed by toxins produced by ComX-dependent cells and lack autoimmunity (9).

In *Myxococcus xanthus*, the nonsporulating cells of the population undergo altruistic PCD during fruiting body formation. Toxin endonuclease MazF and antitoxin protein MrpC compose a toxin-antitoxin (TA) system (24-25). During the formation of fruiting bodies, the modification of the regulatory protein MrpC by cell proteases liberates it from repression (24,26) and causes the activation of hundreds of genes, including itself and the toxin MazF (26-27). The interaction between MazF and MrpC results in the autolysis of nonsporulating cells (24), which ensures the survival of the sporulating subpopulation by providing nutrients.

In all three autolysis mechanisms, regulators play key roles in regulating cell death, i.e., Spo0A in *Bacillus subtilis*, ComE in *Streptococcus pneumoniae* and MrpC in *Myxococcus xanthus*. Genes for toxin synthesis and autoimmunity are either in the same operon (*skf, cib*) or under the control of the same regulator (ComE, MrpC). The toxins leading to cell autolysis are proteins or peptides, i.e., CbpD and MazF are toxic proteins, and SkfA and SdpC are antimicrobial peptides. However, bacterial autolysis is a general differentiation strategy for bacterial social multicellular behavior, the novel factors and mechanisms to trigger autolysis remain scarce.

In this study, cannibalism in the biofilm of a plant beneficial rhizobacterium, *Bacillus velezensis* SQR9, which can colonize plant roots efficiently through biofilm formation and secrete several secondary metabolites protecting the plant from pathogen invasion (28-29), was investigated. Strain SQR9 harbors a novel genomic island GI3, encoding the synthesis of novel branched-chain fatty acids, bacillunoic acids, through a hybrid type I fatty acid synthase (FAS)-polyketide synthase (PKS) system. GI3 also encodes an ABC transporter (*bnaAB* operon) to export toxic bacillunoic acids upon their induction of self-immunity (30). Here, we revealed that bacillunoic acids-induced immunity is mediated by a regulatory protein. The signal produced by bacillunoic acids is transferred to the immunity proteins BnaAB through a two-component system, and this phosphorylation cascade amplification system plays an important role in bacillunoic acids signal transduction. This study revealed a novel non-protein toxin factor and a different regulating mechanism for *Bacillus* cannibalism for enhancing biofilm formation and group competitiveness.

## Results

### Autolysis of the community subfraction promoted the biofilm formation of SQR9

Cell lysis is a mechanism by which toxin-producing cells express killing factors to lyse a fraction of their siblings that have not developed immunity to killing factors. Microscopic examination of SQR9 cells under biofilm-forming conditions revealed that although most of the cells maintained a regular rod shape, some cells were lysed, with broken and punched cells in the biofilm community (Figure 1A), suggesting the possibility that SQR9 undergoes cell lysis during biofilm formation.

**Figure 1.**
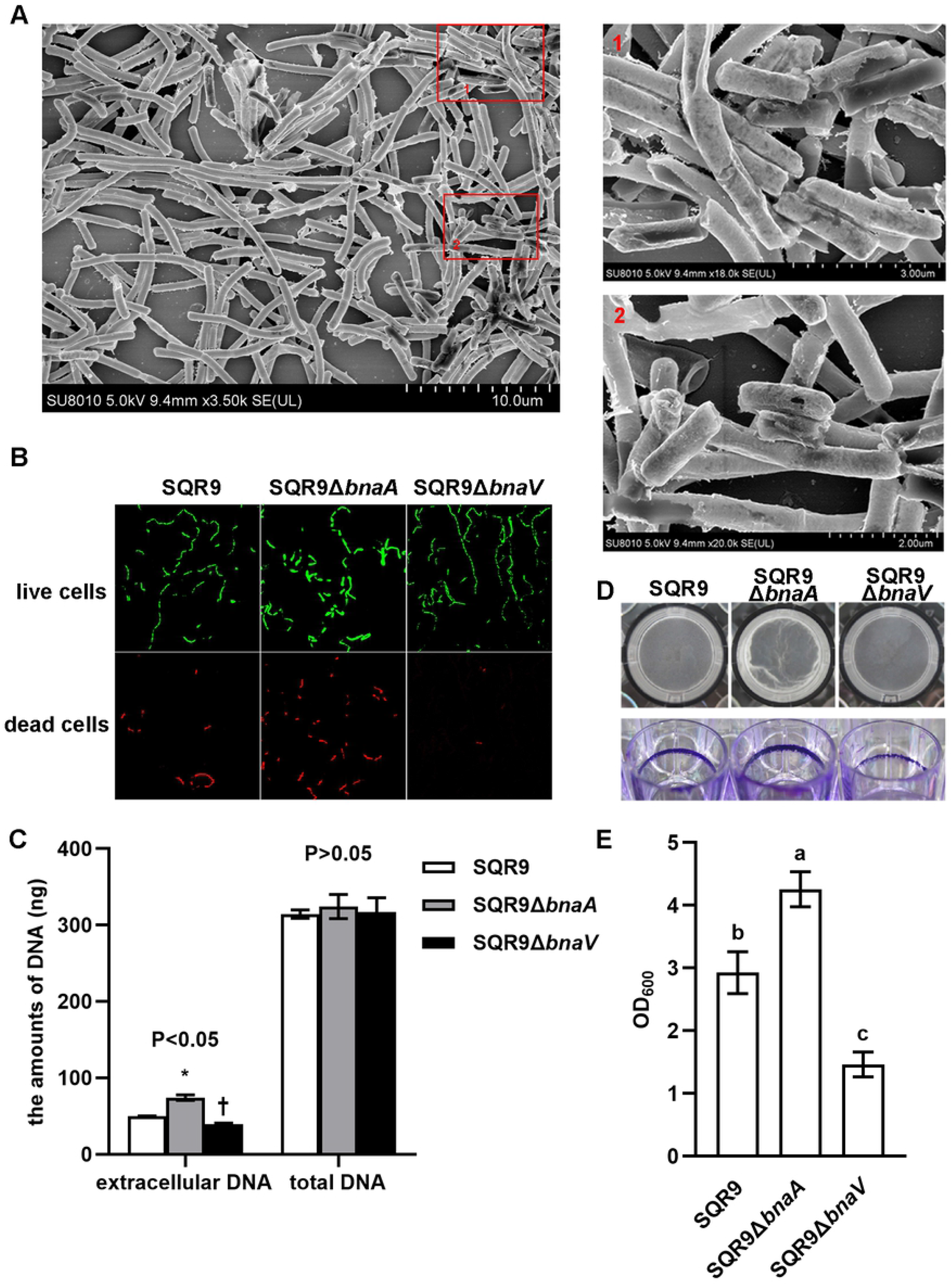
Autolysis of a subfraction of cells promoted the biofilm formation of SQR9. (A) Scanning electron microscopy of cell lysis in SQR9 biofilms. Most of the cells maintained a regular rod shape, but red boxes represent lytic cells (larger versions are shown on the right). (B) Live/dead cell staining of biofilms formed by different strains. Biofilm cells were treated with dyes for live/dead cell staining and examined under CLSM. (C) The amounts of extracellular DNA and total DNA in different strains. * Significant (*p* < 0.05) increase in amounts of extracellular DNA compared with the wild-type SQR9; † significant (*p* < 0.05) decrease in amounts of extracellular DNA compared with the wild-type SQR9. (D) Pellicle formation and the presence of adherent biofilm cells with crystal violet staining. The strains were inoculated into MSgg medium and incubated at 37 °C for 16 h. (E) Quantitative analysis of biofilms with crystal violet staining. OD_600_ values of solubilized crystal violet were from adherent biofilm cells at 16 h. Bars represent standard deviations. Letters above the bars reveal a significant difference based on Duncan’s multiple range test at a *p* < 0.05 level.

To investigate whether the novel genomic island GI3-encoded bacillunoic acids and the ABC transporter plays a role in the biofilm formation of SQR9, the toxin bacillunoic acids synthesis-deficient mutant SQR9Δ*bnaV* and the toxin self-immunity-deficient mutant SQR9Δ*bnaA* (the ABC transporter deletion mutant) (30) were investigated for their biofilm formation. CLSM imaging showed that the dead cells of SQR9Δ*bnaA* were increased and widely distributed around the biofilm (Figure 1B), suggesting that SQR9 in biofilms relies on self-immunity to protect themselves from the lysis of self-produced toxin. However, the dead cells of mutant SQR9Δ*bnaV* were reduced compared to those of wild-type SQR9 (Figure 1B), suggesting that the synthesized bacillunoic acids did play a role in lysing a fraction of their siblings. In the quantitative analysis of cell lysis, extracellular DNA was measured and compared in SQR9, SQR9Δ*bnaA* and SQR9Δ*bnaV*. The amount of total DNA in cultures of these strains appeared to be similar (*p* > 0.05), while compared to wild-type SQR9, the amounts of extracellular DNA of SQR9Δ*bnaA* and SQR9Δ*bnaV* significantly increased and decreased respectively (*p* < 0.05) (Figure 1C).

Biofilm assays showed that SQR9Δ*bnaA* formed stronger while SQR9Δ*bnaV* formed weaker biofilms compared with that of wild-type SQR9 (Figure 1D). Biofilm quantitative analysis (Figure 1E) was consistent with this observation. These results indicated that the cell lysis behavior contributed to biofilm formation.

### Self-immunity during biofilm formation of SQR9 was activated by the toxin

Our previous study suggests that the self-immunity ABC transporter encoded by the *bnaAB* operon was activated by the toxin bacillunoic acids (30). To verify this proposition and explore the induction mechanism, the promoter region of the *bnaAB* operon was fused with the *gfp* reporter gene. The reporter plasmid pNW33N-*P*_*bnaA*_-*gfp* was introduced into SQR9Δ*bnaV* and SQR9 to construct the strains SQR9Δ*bnaV*-*P*_*bnaA*_-*gfp* and SQR9-*P*_*bnaA*_-*gfp*, respectively. During biofilm formation, SQR9-*P*_*bnaA*_-*gfp* showed GFP fluorescence (Figure 2A), but SQR9Δ*bnaV*-*P*_*bnaA*_-*gfp* did not (Figure 2B), suggesting that the *bnaAB* operon could not be expressed in the absence of bacillunoic acids induction. When the fermentation supernatant of wild-type SQR9 was added to MSgg medium, the biofilm cells of SQR9Δ*bnaV*-*P*_*bnaA*_-*gfp* restored GFP fluorescence (Figure 2C), but the fermentation supernatant of the genomic island knockout mutant SQR9ΔGI3 (30) did not have this ability (Figure 2D). These results indicated that the *bnaAB* operon could be activated by bacillunoic acids in the fermentation culture of wild-type SQR9.

**Figure 2.**
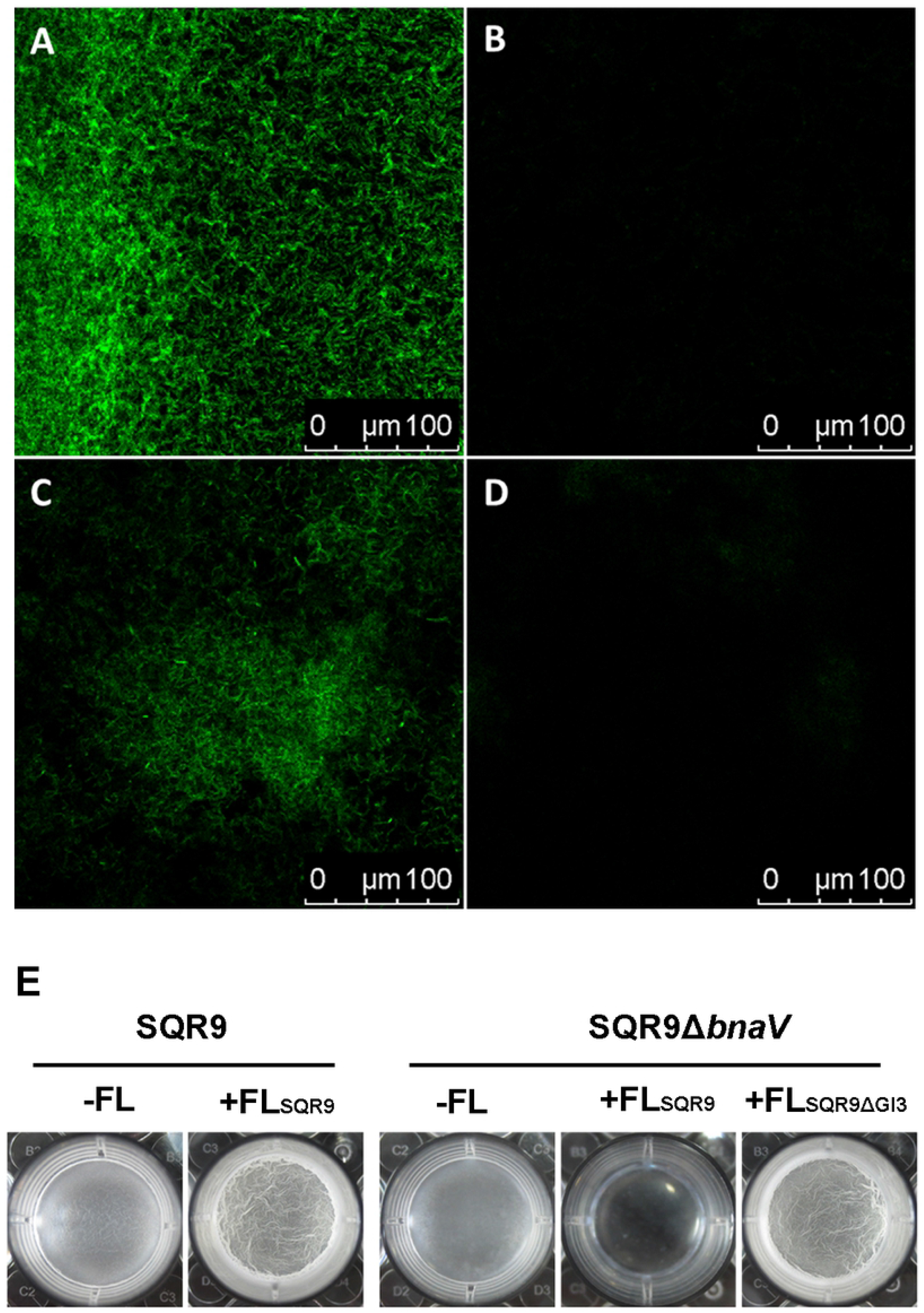
The *bnaAB* operon could be activated by bacillunoic acids. (A-D) GFP fluorescence of strains during biofilm formation: (A) SQR9-*P*_*bnaA*_*-gfp*, (B) SQR9Δ*bnaV*-*P*_*bnaA*_*-gfp*, (C) SQR9Δ*bnaV*-*P*_*bnaA*_*-gfp* with 10% (v/v) of the fermentation liquid of SQR9 and (D) SQR9Δ*bnaV*- *P*_*bnaA*_*-gfp* with 10% (v/v) of the fermentation liquid of SQRMGI3. (E) Comparison of biofilm formation by SQR9 and SQR9Δ*bnaV* in different treatments. The pellicle formed by SQR9Δ*bnaV* with the addition of FL_SQR9_ was very thin at the early stage of biofilm formation.

On the other hand, we observed that the early-stage biofilm formation by SQR9 and SQR9Δ*bnaV* showed some differences in the presence of fermentation liquid (Figure 2E). First, when fermentation liquid of SQR9 (FL_SQR9_) was added to the medium, the SQR9 biofilm was increased and the SQR9Δ*bnaV* biofilm was decreased. Since FL_SQR9_ contains both nutrients and bacillunoic acids, it was speculated that SQR9 biofilm increase was stimulated by the nutrients, whereas bacillunoic acids synthesis defective cells of SQR9Δ*bnaV* were challenged to survive in the presence of exogenous bacillunoic acids in FL_SQR9_. If FL_SQR9_ was replaced by FL_SQR9ΔGI3_ (fermentation liquid of SQR9ΔGI3) that contains nutrients without bacillunoic acids, the defective biofilm formation by SQR9Δ*bnaV* was restored. These results indicated that the wild-type cells with the ability to synthesize bacillunoic acids could actively induce self-immunity against exogenous bacillunoic acids. However, although cells of SQR9Δ*bnaV* could not produce bacillunoic acids, self-immunity of the cells could be induced by the addition of bacillunoic acids, but this induction may not occur timely; only a small number of cells can be successfully induced and survived. In other words, the cells that cannot secrete killing factors and induce self-immunity are very likely to be lysed by the killing factors secreted by toxin-producing cells.

### Cells were differentiated into cannibals and nonproducers during biofilm formation of SQR9

To monitor cell differentiation in biofilm formation, biofilms of strains SQR9-*P*_*bnaA*_-*gfp* and SQR9Δ*bnaV*-*P*_*bnaA*_-*gfp* were observed using CLSM, and cells were stained with PI to distinguish dead cells (dead cells appeared red with PI staining). Only a portion of SQR9-*P*_*bnaA*_-*gfp* cells showed GFP fluorescence (Figure 3), suggesting that at the time point of observation, these cells act as toxin producers, to produce the toxin bacillunoic acids and activate self-immunity ability. Some cells did not show GFP fluorescence (Figure 3), indicating that these cells neither produce bacillunoic acids nor induce self-immunity activity. Some of these cells were dead (appeared red under PI stain) and were probably killed by bacillunoic acids secreted by toxin-producing cells. Strain SQR9Δ*bnaV*-*P*_*bnaA*_-*gfp* lost the ability to produce bacillunoic acids, and no cells showed GFP fluorescence during biofilm development (Figure 3), suggesting the inactivation of the *bnaAB* operon promoter without toxin induction.

**Figure 3.**
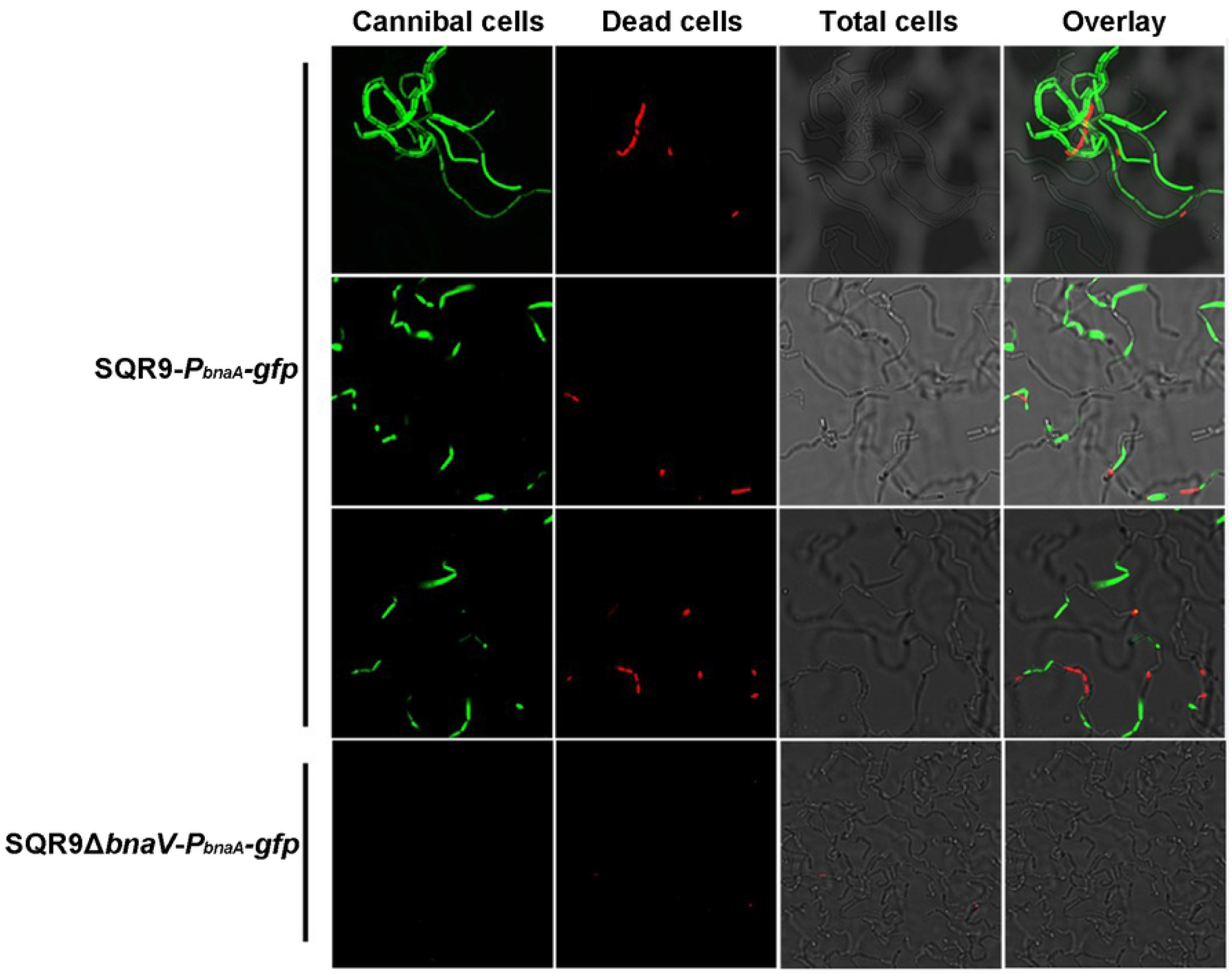
Subpopulations of toxin-producing cells and dead cells were distinct under the microscope. The biofilms of two constructed strains harboring the *P*_*bnaA*_*-gfp* reporter fusion grew on coverslips in MSgg medium. Biofilm cells on coverslips were treated with dyes for dead cell staining, placed on microscope slides and examined under CLSM.

### OmpR and OmpS constitute a two-component system (TCS) to regulate self-immunity expression

The possible regulators controlling the expression of self-immunity ABC transporters were explored, and it has been reported that efflux transporter genes are usually located adjacent to a two-component system for sensing environmental signals in bacteria (31-35). Gene analysis of GI3 showed that the *ompR* gene is located adjacent to the self-immunity ABC transporter gene *bnaAB* (Supplementary Figure S1B). OmpR was predicted to consist of two domains, an N-terminal domain of approximately 111 amino acids joined by a small linker region to a C-terminal domain of approximately 77 residues. The N-terminal domain contains the aspartate residue, which is phosphorylated, and other active-site residues that are highly conserved within the family of response regulator proteins. The C-terminal domain of OmpR is a DNA-binding domain (Supplementary Figure S1C). The heterologously expressed intact OmpR protein and its truncated protein OmpR_c_ (the C-terminal 106 amino acid residues of OmpR) were shown to directly bind to *P*_*bnaA*_ (Figure 4A, 4B), as assessed by biolayer interferometry (BLI), with Kd values of 670±130 nM and 880±140 nM, respectively, at 25 °C. Thus, the protein OmpR regulates the expression of the *bnaAB* operon.

**Figure 4.**
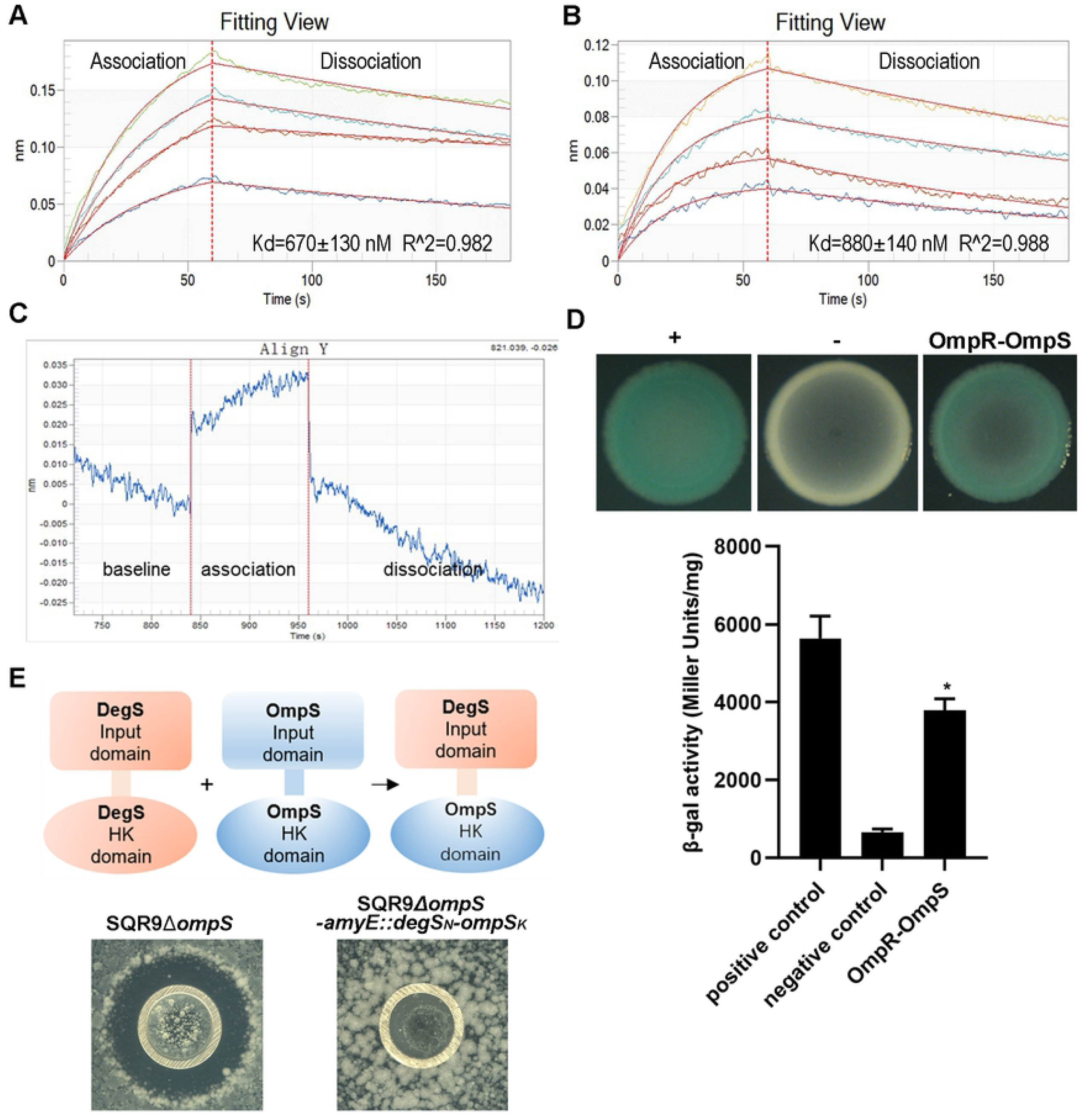
OmpR and OmpS constitute a two-component system (TCS) to regulate self-immunity expression. (A-B) Biolayer interferometry analysis of the interaction between the transcription factor OmpR and *P*_*bnaA*_. Different concentrations of OmpR_c_ (0, 250, 500, 750 and 1000 nM)/OmpR (0, 200, 400, 600 and 1000 nM) were chosen. The figures shown are expanded views of the association and dissociation phases of the sensorgrams: (A) Interaction between *P*_*bnaA*_ and OmpR_c_. (B) Interaction between *P*_*bnaA*_ and OmpR. (C-E) Interaction between OmpS and OmpR. (C) Interaction of OmpS* and OmpR detected by biolayer interferometry (BLI) analysis. (D) Bacterial two-hybrid analysis of OmpR and OmpS protein interactions. The strain containing pKT25-zip and pUT18C-zip was used as a positive control, and the strain containing empty vectors pKT25 and pUT18C was used as a negative control. Cotransformants were spotted onto selective screening LB agar plates and incubated at 30 °C. Blue indicates a positive interaction, and white indicates no interaction. Quantitative analysis of β-galactosidase activity by Miller units (n=3, *p* < 0.05, one-way ANOVA). (E) The kinase domain of OmpS can transfer phosphate groups to OmpR to activate the immunity gene *bnaAB*. A construction diagram of the chimera kinase DegS_N_-OmpS_K_ by recombination with domains of OmpS and DegS is shown in the figure above. Inhibition of the MeOH-extracted SQR9 supernatant on the growth of SQR9Δ*ompS* and SQR9Δ*ompS-amyE::degS*_*N*_*-ompS*_*K*_ is shown in the figure below.

A classical TCS consists of a transmembrane histidine kinase (HK) and a cytoplasmic response regulator (RR) protein. The kinase OmpS and regulatory protein OmpR were predicted to consist of a TCS to control the expression of self-immunity ABC transporters. The interaction between OmpR and the kinase domain of OmpS (designated OmpS*) was evaluated by BLI analysis, which showed a response of interference (Figure 4C). In addition, the interaction between OmpS and OmpR was confirmed using bacterial two-hybrid systems (BACTH), which showed the positive interaction of these two proteins (Figure 4D). Furthermore, we constructed a hybrid sensor kinase to define that OmpS interacts with OmpR only through its kinase domain, regardless of what the input domain is. Inhibition of flagellar rotation acts as a signal to activate histidine sensor kinase DegS (36). The N-terminal domain of OmpS was replaced by the corresponding domain of kinase DegS and resulted in a hybrid kinase DegS_N_-OmpS_K_ (Figure 4E). By hypothesis, self-immunity ability of strain SQR9Δ*ompS-amyE::degS*_*N*_*-ompS*_*K*_ towards bacillunoic acids is controlled by the rotation of its flagella. As expected, SQR9Δ*ompS-amyE::degS*_*N*_*-ompS*_*K*_ cells being plated equally on LB agar could resist bacillunoic acids (in the fermentation liquid of SQR9) (Figure 4E). It could be inferred from this result that inhibition of flagellar rotation acts as a signal to activate the N-terminal domain of DegS, and then the signal is delivered to the OmpS kinase domain, which interacts with OmpR to complete the rest of the immune pathway. These results suggest that the kinase domain of OmpS interacts with OmpR.

### OmpS participates in transmitting signals induced by bacillunoic acids

It is expected that bacillunoic acids activate the autophosphorylation of OmpS, and the phosphorylated OmpS transfers a phosphoryl group to OmpR. To verify this hypothesis, the OmpR phosphorylation level was compared in strain SQR9Δ*ompS*, SQR9Δ*bnaV* and SQR9. Although the total OmpR protein level remained constant (SDS–PAGE), an OmpR mobility shift was observed in wild-type SQR9 but not in SQR9Δ*ompS* or SQR9Δ*banV* (Phos-tagged SDS–PAGE) (Figure 5). The lack of phosphorylation of OmpR in SQR9Δ*ompS* suggests that OmpS and OmpR constitute the TCS mentioned above. The lack of phosphorylation of OmpR in SQR9Δ*banV* indicates the importance of bacillunoic acids in the immune system; bacillunoic acids can activate the autophosphorylation of OmpS, which causes phosphorylation of OmpR. This was also confirmed by the phenomenon in which the phosphorylation level of OmpR in SQR9Δ*bnaV* was increased after the addition of SQR9 fermentation liquid (Supplementary Figure S2). Together, these results suggest that OmpS participates in the signal transduction induced by bacillunoic acids.

**Figure 5.**
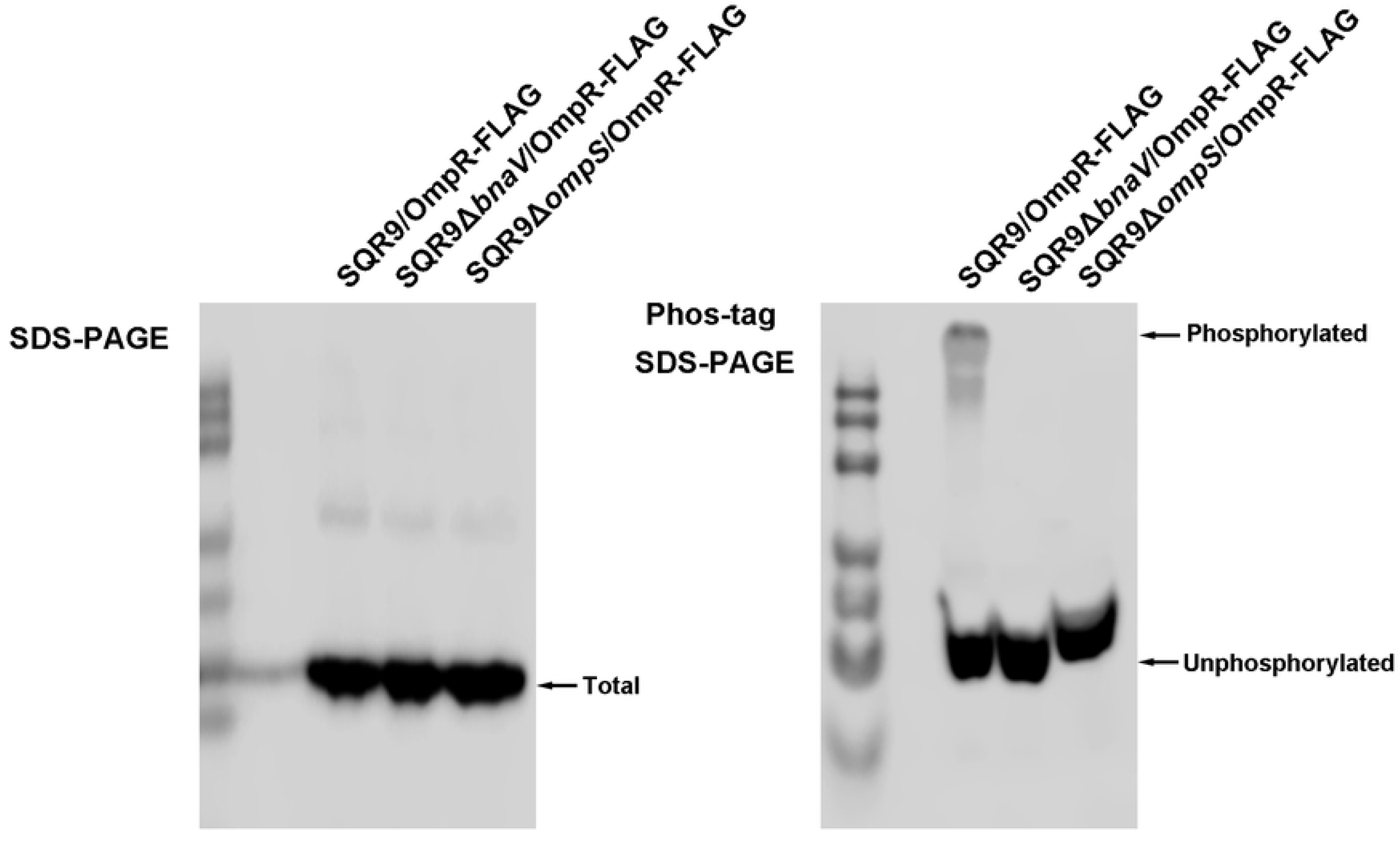
OmpS participates in transmitting signals induced by bacillunoic acids to OmpR. Proteins in the SQR9Δ*ompS*, SQR9Δ*bnaV* and SQR9 strains were separated by normal SDS–PAGE and Zn^2+^- Phos-tag, and OmpR-FLAG was detected by Western blot using the FLAG antibody.

### The GI3-governed cannibalism promoted bacillunoic acids-producing ability of the community

We wonder if the cannibalism-mediated bacterial community optimization to purge the non-producer of the so-called “public goods” can promote the whole community’s competition. We then replaced the bacillunoic acids-inducible promoter with the constitutive promoter *P*_*43*_ to construct a strain SQR9-*P*_*43*_*-bna*, in which self-immunity towards bacillunoic acids is always present regardless of the induction, so non-producing cells in the community cannot be eliminated. Since bacillunoic acids are solely responsible for SQR9 to kill the congeneric strain of *Bacillus velezensis* FZB42 (30), then, the inhibiting abilities of the wild-type SQR9 and non-cannibalism SQR9-*P*_*43*_*-bna* communities against FZB42 were used to indicate the production of bacillunoic acids. Results showed that strain SQR9-*P*_*43*_*-bna* produced less bacillunoic acids than SQR9 did, regardless of the inhibition zone around the spot on the lawn (Figure 6A) or the inhibition zone around the Oxford cup on the plate (Figure 6B), suggesting that cannibalism optimized the bacterial community by eliminating the non-producer of the “public goods” and promoted the whole community’s production and competition.

**Figure 6.**
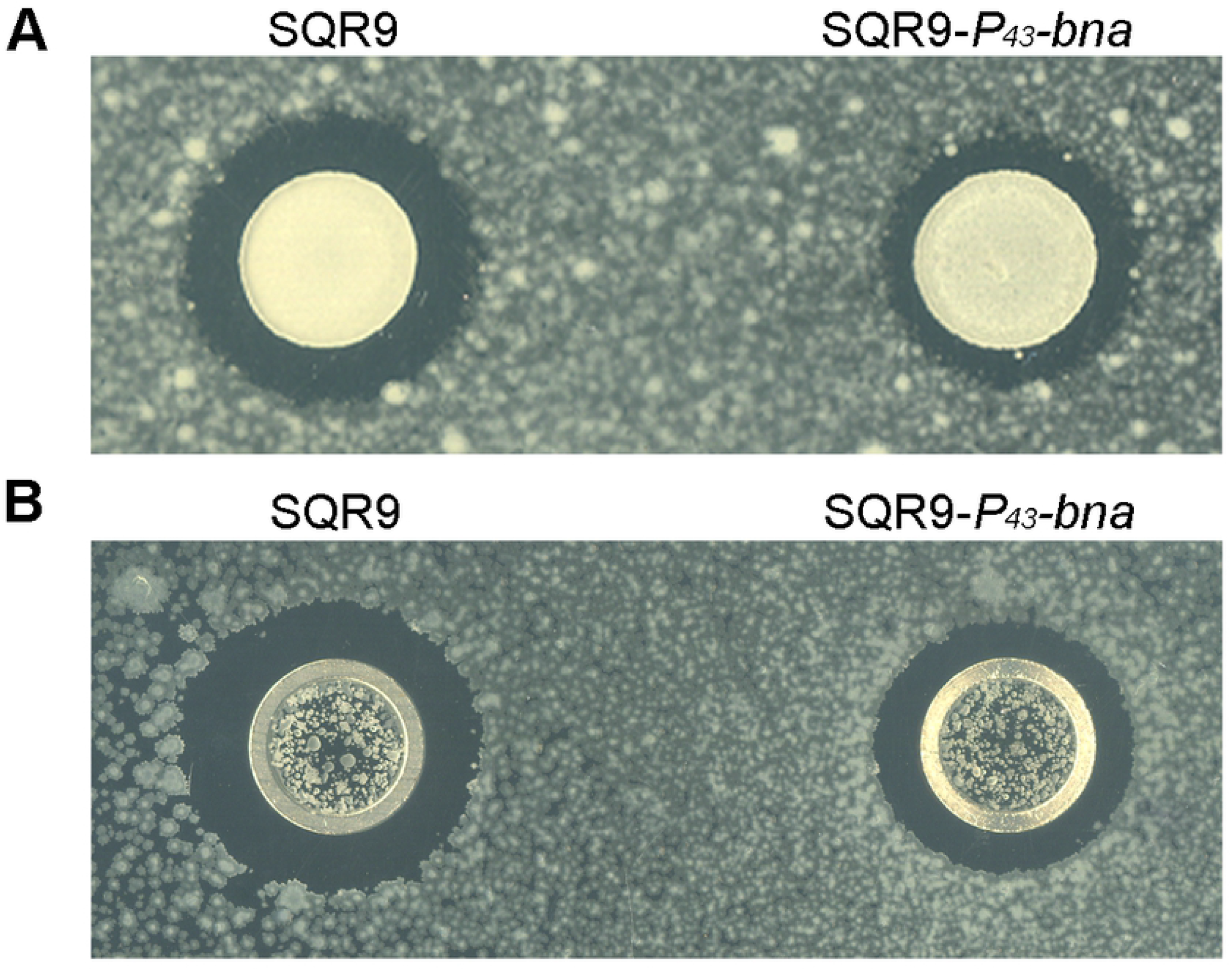
The cannibalism-mediated bacterial community optimization promoted the community’s competition. The inhibiting abilities of the wild-type SQR9 and non-cannibalism SQR9-*P*_*43*_*-bna* communities against strain FZB42 were evaluated with (A) colony and (B) Oxford cup inhibition on the plate.

## Discussion

We have elucidated a genomic island (GI3)-governed cannibalism phenomenon in *B. velezensis* SQR9 biofilm formation, which involves the secondary metabolic toxins bacillunoic acids; a two-component signal-transduction system, OmpS-OmpR; and an ABC transporter for self-immunity, BnaAB. The work presented here demonstrates that a subpopulation of cells within the biofilm differentiate into cannibals and secrete toxic bacillunoic acids. When the presence of bacillunoic acids activates the autophosphorylation of kinase OmpS in cells of this subpopulation, it transfers a phosphoryl group to its response regulator OmpR, the phosphorylation of which induces the transcription of ABC transporter BnaAB to pump bacillunoic acids out of the cell for self-protection. The exported toxins could lyse the non-producers that lack this self-immunity, and the release of nutrients to promote biofilm formation (Figure 7).

**Figure 7.**
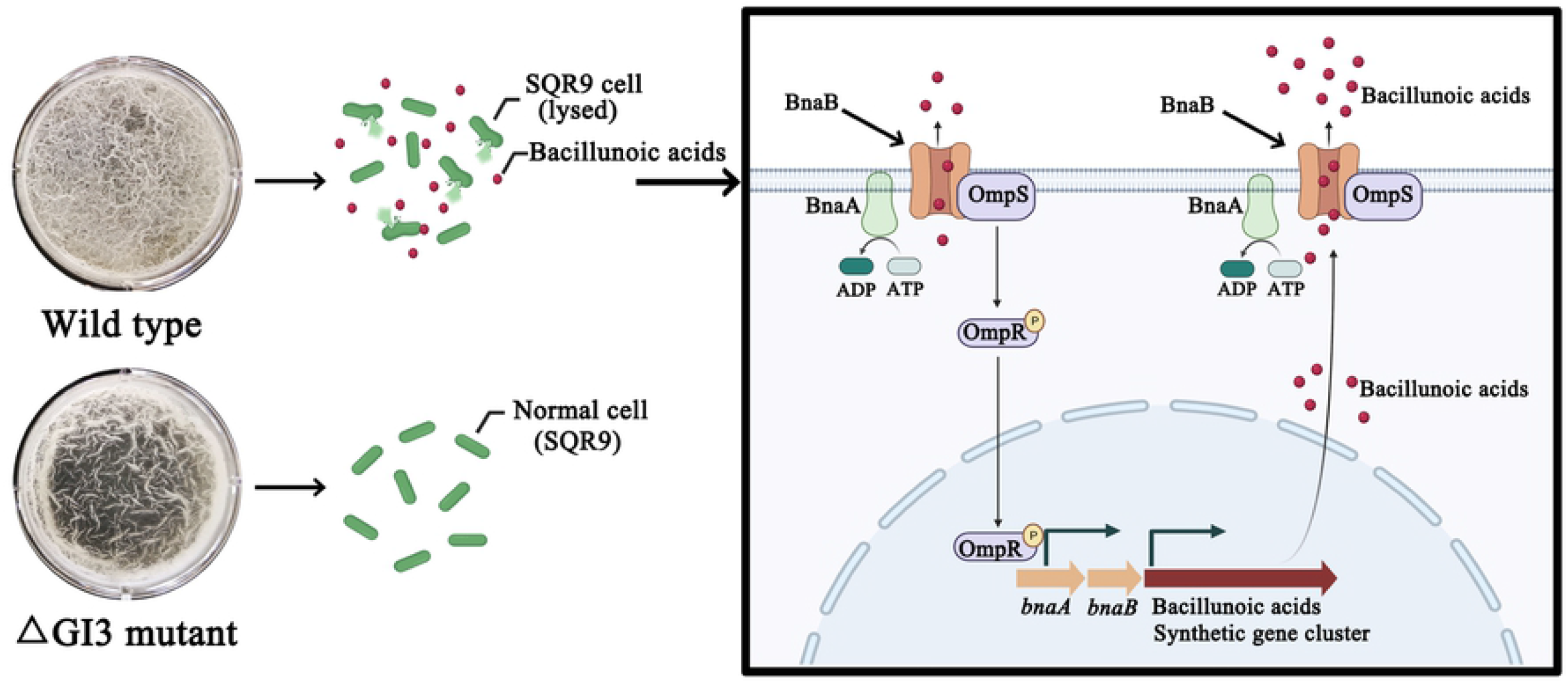
Genomic island (GI3)-governed cannibalism contributes to biofilm formation in *B. velezensis* SQR9. In this model, a subpopulation of cells within the biofilm differentiates into cannibals and secretes toxic bacillunoic acids. When the presence of bacillunoic acids activates the autophosphorylation of kinase OmpS in cells of this subpopulation, it transfers a phosphoryl group to its response regulator OmpR, the phosphorylation of which induces the transcription of ABC transporter BnaAB to pump bacillunoic acids out of the cell for self-protection. Exported toxins can lyse nonproducers that lack self-immunity, and the release of nutrients such as eDNA promotes biofilm formation.

In the process of multicellular differentiation, the killing of genetically identical cells for the benefit of the community is an important phenomenon. Usually, this phenomenon is based on the molecular mechanisms of killing and immunity to toxins (37). To date, although three different strategies for controlling the death and life of bacterial communities have been widely studied in distinct model organisms, they might have some common features. For example, among the three mechanisms of cannibalism (*B. subtilis*), fratricide (*S. pneumoniae*), and altruistic autolysis (*M. xanthus*), the self-immunity genes and toxin-synthesizing genes are generally in the same operon (*skf, cib*) or controlled by the same regulator (ComE, MrpC) (10,13,26,38). Only the three-protein signal-transduction cannibalism system of *B. subtilis* is an exception, in which the toxin SdpC induces the immune system by acting as a ligand that binds to the signal-transduction protein SdpI and subsequently sequesters the autorepressor SdpR (15). In our study, self-immunity genes and toxin-synthesizing genes involved in cannibalism were encoded by a novel genomic island GI3, which has never been reported before. Similar to the behavior of toxin SdpC in *B. subtilis*, toxin bacillunoic acids also induce the immune system (Figure 2). However, this induction is achieved in a completely different way. The *bnaAB* gene for immunity against bacillunoic acids was activated by the two-component system (TCS) OmpS-OmpR, which is regulated by phosphorylation. Given that TCS constitutes phosphoryl transfer pathways, it has the characteristics of signal amplification (39). We propose that even a small amount of bacillunoic acids is produced, the induction of self-immunity in cannibalism of SQR9 will still be precise and effective because of signal amplification by TCS.

The phenomenon that a TCS (OmpS-OmpR) governs the ABC transporter (BnaAB) for self-immunity is not unusual. Generally, bacteria in the environment have developed various mechanisms to resist antimicrobial agents. As one of these strategies, regulation of the detoxification mechanism by TCS-controlled efflux pumps is a common mode. For example, in *Bacillus subtilis*, the TCS (BceRS) induces the expression of the bacitracin transporter BceAB (40). In *Staphylococcus aureus*, two ABC transporters, BraDE and VraDE, regulated by the TCS (BraRS), are involved in bacitracin and nisin resistance, respectively (41). In *Bacillus licheniformis*, BacRS acts as a TCS to control the expression of the bacitracin transporter gene *bcrABC* (33). However, pumping out of these toxins by the transporter are not involved with the killing of genetically identical cells mentioned in this study. In contrast, we discovered that in the bacillunoic acids synthesis mutant SQR9Δ*bnaV*, the proportion of cell lysis and the amounts of extracellular DNA and biofilm formation were decreased compared with those of wild-type SQR9 (Figure 1). These results indicated that *B. velezensis* SQR9 used bacillunoic acids as toxins to lyse part of their sibling cells for the benefit of the population. On the other hand, bacterial cannibalism toxins in previous studies are usually proteins or peptides that are produced through ribosomal synthesis pathways, such as toxin peptides SdpC and SkfA in *B. subtilis* (42); toxin proteins CbpD and LytA in *S. pneumoniae* (22,43) and toxin protein MazF in *M. xanthus* (24). However, the toxins bacillunoic acids are secondary metabolites synthesized by the nonribosomal pathway in SQR9, which have favorable features of high thermostability, photostability and pH stability in the external environment (30).

The mechanism of bacillunoic acids sensing remains unknown, we tried to construct a hybrid kinase to determine whether the N-terminus of OmpS could sense bacillunoic acids. To indicate the sensing of bacillunoic acids by OmpS, its kinase domain was replaced by that of DegS, resulting in the hybrid kinase OmpS_N_-DegS_K_ (Supplementary Figure S3A). Since the DegSU system of Bacillus is well-known to regulate the biofilm formation, swarming motility and complex colony architecture (44), once the hybrid kinase OmpS_N_-DegS_K_ senses the bacillunoic acids, then the phosphorylated DegU will activate genes involved in biofilm formation. It was expected that the biofilm formation ability of strain SQR9Δ*degS-amyE::ompS*_*N*_*-degS*_*K*_ was controlled by bacillunoic acids. However, this strain did not exhibit a three-dimensional pellicle biofilm structure similar to wild-type SQR9 (Supplementary Figure S3B), possibly due to that the construction of this hybrid kinase could be problematic, or the N-terminal domain of the chimera kinase OmpS_N_-DegS_K_ may not be able to sense bacillunoic acids and therefore does not transmit the signal to downstream DegSU. Detailed analysis showed that only 3 amino acids at the N-terminus of OmpS were outside the membrane (Supplementary Figure S4), which is too short and can only be buried inside the cytoplasmic membrane. According to reports, the N-terminal extracellular domain of intramembrane-sensing histidine kinases (IMHKs) is usually too short to perceive stimuli directly; thus, minimalistic HKs need to recruit accessory membrane proteins as sensors (45-47). For example, in *B. subtilis*, IMHK BceS does not sense bacitracin directly but forms a sensory complex with the permease BceB, which recognizes bacitracin and acts as the stimulus for BceS activation (48). Thus, OmpS is probably an IMHK, and the N-terminal domain of OmpS did not act as a sensor and relied on an accessory membrane protein to sense the toxin bacillunoic acids. To search for the possible accessory protein of OmpS, bacterial two-hybrid systems were used to identify protein–protein interactions between OmpS and other proteins encoded by the genes adjacent to *ompS* in GI3, no interaction between OmpS and the ATPase BnaA was observed, but OmpS clearly interacted with the permease BnaB. In addition, the OmpS-OmpS pair showed positive interaction results, forming a dimer of the kinase as expected (Supplementary Figure S5). Thus, OmpS may form a sensory complex in the membrane with permease BnaB, which acts as an accessory protein to sense bacillunoic acids and transfer the signal to OmpS and the downstream response regulator OmpR. But the combination of BnaB and OmpS needs further investigation.

The two antimicrobial peptides, SkfA and SdpC, which are associated with cannibalism, are produced almost exclusively by *Bacillus subtilis* and are hardly found in other microorganisms. Specifically, except for *B. subtilis*, the *skf* locus is only found in *Paenibacillus larvae*, and is not found outside the genus; the *sdpABC* operon is only preserved in *Bacillus clausii* and the *Bacillus cereus* plasmid pBC239 (9). In our study, the autolysis element of *B. velezensis* SQR9 is located on the genomic island GI3, which is speculated to be obtained from other microorganisms by horizontal gene transfer (30). It is indicated that similar autolysis elements and cannibalism phenomena might exist in microorganisms other than *B. velezensis* SQR9.

Bacillunoic acids are the major compounds for the competition with closely related bacteria (30). We discovered that SQR9 produced more bacillunoic acids in the presence of FZB42 than the SQR9 monoculture, regardless of the indirect interaction of fermentation liquid or direct cell contact. The amount of bacillunoic acids production was indicated by the inhibition zone around the Oxford cup on the plate (Supplementary Figure S6A), which was produced only by the antagonistic activity of bacillunoic acids against FZB42, as reported by Wang et al. (30). In addition, as mentioned above, the *bnaAB* operon was activated by bacillunoic acids, and the strength of *P*_*bna*_ in strain SQR9-*P*_*bna*_*-gfp* under different treatments was also tested and indicated by the green fluorescence intensity. The interaction of SQR9 with FZB42 promoted stronger *P*_*bna*_ ability than single-strain SQR9 (Supplementary Figure S6B). This result suggested that SQR9 takes advantage of this toxin to suppress the growth of phylogenetically closed bacteria and compete for the same resources. Based on this observation, we proposed an additional role for bacillunoic acids in protecting cells within the biofilm against competing *Bacillus* relatives.

In the environment, SQR9 and its homologous strains have the same living habit. Once food shortages occur, they are forced to adjust their communities to adapt to the environment and competewith each other for survival. Overall, the genomic island GI3 is more like an excellent weapon for SQR9, mainly reflected in two aspects. On the one hand, GI3 mediated the killing of the sibling cells involved in cannibalism within the community, which were scarified for the benefit of the whole population. On the other hand, GI3 also mediated the killing of other organisms when enemies appeared, which ensured the stability of the *B. velezensis* SQR9 community. Whether in intraspecies differentiation and interspecies competition, GI3 is responsible not only for killing but also for self-protection. This is probably an important reason why GI3 was preserved by *B. velezensis* SQR9 during evolution.

## Materials and methods

### Strains, plasmids and growth conditions

The bacterial strains and plasmids used in this study are listed in Supplementary Table S1. *Bacillus velezensis* SQR9 (CGMCC accession No. 5808; China General Microbiology Culture Collection Center; NCBI accession No. CP006890) was used throughout this study. *Bacillus velezensis* FZB42 (49) was used to test the bacillunoic acids-producing ability of wild-type SQR9 and its mutants. *E. coli* TOP 10 (Invitrogen, Shanghai, China) was used as the host for all plasmids. *Bacillus* strains and *E. coli* TOP 10 were routinely grown at 37 °C in Luria-Bertani (LB) medium (peptone, 10 g/L; NaCl, 3 g/L; yeast extract, 5 g/L). For biofilm formation, *B. velezensis* SQR9 and its mutants were cultivated for 16 h in MSgg medium (50) at 37 °C. For secondary metabolite production and fermentation liquid collection, *B. velezensis* SQR9 and SQR9ΔGI3 were grown for 48 h in Landy medium (51) containing 20 g/L glucose and 1 g/L yeast extract at 30 °C. When necessary, antibiotics were added to the media at the following final concentrations: zeocin, 20 μg/mL; spectinomycin, 100 μg/mL; kanamycin, 30 μg/mL; ampicillin, 100 μg/mL; chloramphenicol, 5 μg/mL for *B. velezensis* strains and 12.5 μg/mL for *E. coli* strains; erythromycin, 1 μg/mL for *B. velezensis* strains and 200 μg/mL for *E. coli* strains. The media were solidified with 2.0% agar.

### Gene knockout by allelic exchange

To delete the target genes, the upstream and downstream regions that flanked the target genes were amplified from the SQR9 genome, and the erythromycin resistance gene was amplified from plasmid pAX01. The fragments were fused through overlap PCR, the fused fragments were transformed into competent cells of *B. velezensis* SQR9, and the transformants were screened and verified as described in a previous study (52). The primers used in this study are shown in Supplementary Table S2.

### Construction of *P*_*bnaA*_-*gfp* expression fusions in *B. velezensis* strains

The promoter-less *gfp* gene has been integrated into *B. velezensis*-*E. coli* shuttle vector pNW33N to generate the plasmid pNW33N-*gfp* (53). The *P*_*bnaAB*_ fragment was used to construct pNW33N-*gfp*. In detail, the promoter fragment bordered by primer-introduced *Pst1* and *BamH1* sites was amplified from SQR9 genomic DNA using the primers Pst1-F and BamH1-R. After purification and digestion with the corresponding restriction enzymes, the fragment was cloned into plasmid pNW33N-*gfp*, resulting in plasmid pNW33N-*P*_*bnaA*_-*gfp*. This plasmid was transformed into SQR9 and SQR9Δ*bnaV*. The primers used in this study are shown in Supplementary Table S2.

### Biofilm assay and microscopic observation

The suspensions of *B. velezensis* strains were prepared and inoculated into the wells of 24-well polyvinylchloride (PVC) microtiter plates (Fisher Scientific, Shanghai, China) filled with MSgg medium as described previously (52). Subsequently, 14-by 14-mm sterile coverslips were inserted into the above wells, and the microtiter plates were incubated at 37 °C without shaking. For scanning electron microscopy, coverslips with biofilm samples were fixed with 2.5% glutaraldehyde for 24 h at 4 °C. Then, the samples were dehydrated, freeze dried, sputter-coated with gold-palladium and visually inspected under a scanning electron microscope (SU8010, Hitachi, Japan). For the visualization of live and dead cells, coverslips with biofilm samples were stained by using a LIVE/DEAD BacLight Bacterial Viability Kit (L7012, Molecular Probes, Invitrogen). Two stock solutions of stain (propidium iodide and SYTO 9) were each diluted to a concentration of 3 μL/mL. Propidium iodide-stained dead cells and SYTO 9-stained live cells in biofilms were visualized by using confocal laser scanning microscopy (TCS SP8, Leica, Germany) with an excitation wavelength of 485 nm, and fluorescence was recorded at wavelengths of 530 nm (emission 1; green) and 630 nm (emission 2; red). When required, 10% (v/v) of the fermentation liquid of SQR9 or SQR9ΔGI3 was added to the MSgg medium. Quantification of biofilms with crystal violet (CV) staining was carried out as described previously (52).

### Extraction and quantification of total DNA and extracellular DNA

DNA was extracted and measured by using a modified method described by a general protocol. The strains were grown overnight at 37 °C in LB medium. The amount of extracellular DNA (eDNA) released from the lysed cells was obtained from the culture supernatant. The cells in culture were removed by centrifugation, and then 3 M sodium acetate and 100% ice-cold ethanol were added to the culture supernatant. The samples were mixed well and stored at -20 °C for 1 h. After centrifugation, the pellet was resuspended in 70% ice-cold ethanol. Next, the ethanol was removed, and the pellet was resuspended in 100 μL of TE buffer. The amount of total DNA was obtained by lysing all cells by sonication to release DNA, and the cell debris was removed by centrifugation. The supernatant was then transferred to a new centrifuge tube, and the same protocol as that for ethanol precipitation of eDNA was performed.

The amount of DNA was quantified by qPCR using the single-copy housekeeping gene *recA*. Serial dilution of genomic DNA was used to generate the standard curve. qPCR analysis was performed using a SYBR Premix Ex Taq kit (TaKaRa, Biotek, Dalian) and an ABI 7500 system under the following conditions: initial cycle at 95 °C for 10 s and then 40 cycles at 95 °C for 5 s and 60 °C for 34 s. The primers used in this study are shown in Supplementary Table S2.

### Cloning, heterologous overexpression and purification of OmpR, its C-terminal fragment (OmpR_c_) and a truncated form of OmpS (OmpS*)

The gene segment encoding the kinase domain of protein OmpS without the terminal transmembrane amino acid sequence was PCR amplified from the genomic DNA of SQR9 with primers pMAL-*ompS\**F and pMAL-*ompS\**R (76Ile-298Gly). Likewise, the full-length gene of OmpR was amplified with primers pMAL-*ompR*-*his*-F and pMAL-*ompR*-*his*-R. The resulting fragments were cloned into an expression vector, pMAL-c5X, to generate the N-terminal MBP (maltose-binding protein)-tagged fusion proteins of OmpS* and OmpR. The DNA fragments encoding full-length OmpR in the vector pMAL-c5X had an additional HHHHHH tag at the C-terminus for removal of the MBP tag with a Ni column. In addition, the DNA fragment encoding OmpR_c_ (the DNA-binding domain of OmpR, Asn121-Val226) was amplified with primers pET-ompR_c_F and pET-ompR_c_R using SQR9 genomic DNA as a template. The PCR product was cloned into the expression plasmid pET-29a (+) with a C-terminal His tag. All constructed plasmids were confirmed by DNA sequencing. The resulting plasmids were individually introduced into *Escherichia coli* BL21(DE3).

*Escherichia coli* BL21(DE3) harboring the expression plasmids pET-29a (+)-*ompR*_*c*_, pMAL-c5X-*ompR* and pMAL-c5X-*ompS\** were grown at 37 °C with shaking at 170 rpm in 100 mL of LB medium containing different antibiotics (kanamycin, ampicillin and ampicillin). When the cell concentration reached an OD_600_ of approximately 0.5, the growth temperature was lowered to 16 °C. After another 30 min of incubation, isopropyl-β-d-thiogalactoside (IPTG) was added to a final concentration of 24 mg/mL to induce the expression of the target proteins. After cultivation at 16 °C with 120 rpm shaking for approximately 12 h, the cells were harvested by centrifugation at 11000×g for 10 min and stored at - 80 °C until use. Collected cells were thawed at room temperature, washed at least twice with 0.01 M phosphate-buffered saline (PBS, 10 mM Na_2_HPO_4_, 138 mM NaCl, 1.8 mM KH_2_PO_4_, 2.7 mM KCl, pH 7.4) and finally resuspended in 20 mL of the same buffer. Thereafter, the suspension was lysed by sonication (3 s pulse on, 3 s pulse off) in an ice bath for 20 min and then centrifuged at 11,000×g for 50 min at 4 °C.

The lysates were passed through a 0.22 μm filter (Millipore) to remove any other aggregates or insoluble particles. The His-tagged OmpR_c_ protein was produced as a soluble protein in *E. coli* BL21(DE3) cells, purified by adsorption to a Profinity Ni-charged IMAC (Bio–Rad) column and eluted stepwise with imidazole. Ni column-bound His-tagged protein was washed with wash buffer containing 30 mM imidazole, and the protein was eluted with elution buffer containing 300 mM imidazole. OmpR and OmpS* were purified by amylose resin (NEB) according to the manufacturer’s instructions. In addition, intact OmpR fused with the MBP tag was purified with a Ni affinity column to remove the tag. After the addition of loading buffer, the prepared samples were incubated for 10 min at 95 °C. After confirmation with 10% sodium dodecyl sulfate polyacrylamide gel electrophoresis (SDS–PAGE), the purified proteins were dialyzed against PBS buffer, concentrated by using an Amicon Ultra15 (MW 10000; Millipore) and stored at -80 °C until further use.

### Biolayer interferometry (BLI) measurements

To confirm whether OmpR can bind *P*_*bnaAB*_ directly, BLI analysis, which can indicate the interaction between protein and DNA fragment, was performed. Samples or kinetics buffers (PBS, PBST: PBS containing 0.02% Tween-20) were dispensed into 96-well black microtiter plates (Millipore, Billerica, MA) with 200 μL in each well. Streptavidin sensor tips were prewetted in PBS for 10 minutes. Determination of binding kinetics was performed on an Octet-RED 96 device at 25 °C with orbital sensor agitation at 1000 rpm. First, biotin-labeled *P*_*bnaAB*_ was immobilized on the biosensor tips at a concentration of 100 nM for 600 s, and a negative control was included using biosensor tips immobilized with PBS. Then, a baseline measurement was performed in the buffer for 300 s. Association was performed using increasing concentrations of OmpR_c_/OmpR (MBP fusion) for 600 s. Subsequently, the biosensor tips were agitated in buffer for 10 min for disassociation. The binding profile of each sample was summarized as the “nm shift” (the wavelength/spectral shift in nanometers), which represents the association and disassociation of the protein and the potential ligands. Sensorgrams were fitted using a 1:1 Langmuir binding model (Analysis Software version 9.0) after subtraction of the control curve date (without ligand) (54).

### Bacterial two-hybrid assay

The Bacterial Adenylate Cyclase Two-Hybrid System (BACTH System Kit, Euromedex, France) was used to analyze protein–protein interactions. The OmpR and OmpS gene sequences were amplified from the *B. velezensis* SQR9 genome and cloned in frame into the expression vectors pUT18C and pKT25, respectively. Screening of the constructed plasmid was performed by sequencing, and the two plasmids were cotransformed into *E. coli* strain BTH101, which has a *lacZ* gene under the regulation of a cAMP-inducible promoter. When the protein physical interaction occurs, the T25 and T18 fragments of adenylate cyclase produce an active enzyme, resulting in synthesis of cAMP and subsequent expression of the *lacZ* reporter gene, as described previously by Karimova et al. (55). Cotransformants were spotted onto LB agar plates containing kanamycin, ampicillin, X-gal (40 μg/mL) and IPTG (0.5 mM). Pictures were taken after 48 h of growth at 30 °C. Negative control transformants (with empty vectors pKT25 and pUT18C) and positive control transformants (pKT25-zip and pUT18C-zip) were included.

The BACTH System was also used to test interactions between proteins encoded by genes of the first operon of the GI3 island. The *ompS* gene was cloned into the vector pKT25, and then *ompR, bnaA, bnaB* and *ompS* were cloned into the vector pUT18C. Primers used were listed in Supplementary Table S2.

### β-galactosidase activity

The efficiencies of interactions between proteins were quantified by measuring β-galactosidase activity in liquid cultures. For this measurement, the transformants were cultivated with vigorous agitation for 24 h at 30 °C in LB medium supplemented with kanamycin, ampicillin and IPTG. The cultures were diluted 1:5 into M63 medium, and the optical density at 600 nm (OD_600_) was recorded. Cells were permeabilized using toluene and 0.1% (w/v) SDS. The tubes were subjected to vortexing for 10 s and incubated at 37 °C for 30 to 40 min to evaporate toluene. For the enzymatic reaction, the β-galactosidase activity was tested (56). In detail, aliquots (0.1 to 0.5 mL) of permeabilized cells were added to buffer PM2 (70 mM Na_2_HPO_4_, 30 mM NaHPO_4_, 1 mM MgSO_4_, and 0.2 mM MnSO_4_, pH 7.0) containing 100 mM β-mercaptoethanol to a final volume of 1 mL. The tubes were incubated at 28 °C in a water bath for 5 min. The reaction was started by adding 0.25 mL of 0.4% ONPG in PM2 buffer (without β- mercaptoethanol). The reaction was stopped by adding 0.5 mL of a 1 M Na_2_CO_3_ solution, and the OD_420_ was then recorded. The enzymatic activity, A (in units per milliliter), was calculated according to the following equation: A = 200 × (OD_420_ of the culture - OD_420_ in the control tube)/minutes of incubation) × dilution factor. One unit of β-galactosidase activity corresponds to the hydrolization of 1 nmol of ONPG per min at 28 °C. The specific activity of β-galactosidase is defined in units per milligram (dry weight) of bacteria. The OD_600_ of the bacterial culture was determined by considering that 1 mL of culture at an OD_600_ of 1 corresponds to 300 μg (dry weight) of bacteria.

### Chimera kinases assay

The first chimera kinase was created by connecting the sensing domain of protein DegS to the kinase domain of OmpS. This fusion segment was then introduced into the *amyE* site of the SQR9Δ*ompS* mutant by homologous flanking fragments. This strain was constructed to address the hypothesis that the phosphorylated kinase domain of OmpS can transfer the phosphoryl group to OmpR and then activate the self-immunity.

Another chimera kinase was created by connecting the sensing domain of protein OmpS to the kinase domain of DegS. The fusion segment was then introduced into the *amyE* site of the SQR9Δ*degS* mutant by homologous flanking fragments. This strain was constructed to address the hypothesis that the N-terminus of OmpS can sense bacillunoic acids. The primers used in the chimera kinase assay are listed in Supplementary Table S2.

### Phosphorylation analysis of OmpR by Phos-tag gel electrophoresis

The *ompR-flag* fusion was inserted into the plasmid pNW33N by the primers listed in Supplementary Table S2, and the recombinant plasmid was transformed into wild-type SQR9, SQR9Δ*ompS* and SQR9Δ*bnaV*. These three strains were incubated at 37 °C and 170 rpm for total protein extraction. Relative OmpR phosphorylation levels were measured with Phos-tag™-acrylamide gel electrophoresis and Western immunoblotting. Samples were resolved on 8% SDS-polyacrylamide gels prepared with 50 mM Phos-tagged acrylamide (WAKO) and 100 mM ZnCl_2_ as described by the manufacturer’s instructions. Before transferring, gels were equilibrated in transfer buffer with 10 mM EDTA for 20 min two times, followed by transfer buffer without EDTA for another 20 min. Protein was transferred from the Phos-tag acrylamide gel to the PVDF membrane for 30 min at 380 mA at 4 °C, and then the membrane was analyzed by western blotting using the anti-FLAG antibody.

### Bacillunoic acids-producing ability of strain SQR9 and the cannibalism mutant communities

The bacillunoic acids-inducible promoter of SQR9 was replaced with the constitutive promoter *P*_*43*_ to construct a strain SQR9-*P*_*43*_*-bna* without cannibalism ability. The primers used in this study are shown in Supplementary Table S2.

For spot-on-lawn assay, the diluted bacterial suspension of FZB42 was spread on an LB plate and used as the lawn after drying. Strain SQR9 and SQR9-*P*_*43*_*-bna* were grown at 37 °C with shaking until stationary phase. The cultures were adjusted to the same OD_600_ and spotted with a volume of 2 μl on the FZB42 lawn on each plate. The plates were incubated for 36 h at 22 °C, and the antibacterial activity was observed as a clear zone around the spot.

For Oxford cup assay, at the stationary phase, bacterial suspension of SQR9 and SQR9-*P*_*43*_*-bna* were adjusted to the same OD_600_ and harvested by centrifugation at 10 000 × g for 10 min. The supernatant was dehydrated by a freeze-dryer and then extracted by methanol (5 mL of MeOH for 20 mL of supernatant). The diluted bacterial suspension of FZB42 was spread on an LB plate and used as the lawn after drying. An Oxford cup with a volume of 100 μL of MeOH extraction was placed on the plate. The plates were incubated for 36 h at 22 °C, and the antibacterial activity was observed as a clear zone around the Oxford cup.

### Bacillunoic acids induction experiment

Strains of SQR9 and FZB42 were grown overnight at 37 °C with shaking. They were premixed at a 1:1 ratio based on their OD_600_ values, and the mixture was inoculated into fresh LB medium at 1%. At the same time, a single-strain culture of SQR9 was inoculated into fresh LB medium at 1%. When the OD_600_ was recorded as 0.5, fermentation liquid of FZB42 was added at 10% of the total volume. At the stationary phase, each bacterial suspension was adjusted to the same OD_600_ and harvested by centrifugation at 10 000 × g for 10 min. The supernatant was dehydrated by a freeze-dryer and then extracted by methanol (5 mL of MeOH for 20 mL of supernatant). The diluted bacterial suspension of FZB42 was spread on an LB plate and dried. An Oxford cup with a volume of 100 μL of MeOH extraction was placed on the plate. The plates were incubated for 36 h at 22 °C, and the antibacterial activity was observed as a clear zone around the Oxford cup.

Stationary phase bacterial cultures of SQR9-*P*_*bnaA*_*-gfp*, SQR9-*P*_*bnaA*_*-gfp* with fermentation liquid of FZB42, and SQR9-*P*_*bnaA*_*-gfp* with FZB42 were acquired as mentioned above. Two hundred microliters of each culture were applied to measure the OD_600_ and green fluorescence intensity, excited at 488 nm and measured at 533 nm using a fluorescence microplate reader.

## Acknowledgements

This research was financially supported by the National Natural Science Foundation of China (31870096).

## Author Contributions

R.Z., Z.X., Q.S. and N.Z. contributed to the conception and design of the study. J.S. provided all reagents used in this study. R.H. and Q.L. performed the experiments. H.F. gave a guide to the BLI experiments. R.H., Q.L., D.W. and N.Z. interpreted the results. R.H. and Q.L. and D.W. prepared the manuscript. Z.X. drew a model graph of the results. R.Z. and Z.X. discussed the results and revised the manuscript.

